# Small form factor implantable neural probe with efficient flip chip μLED for in vivo optogenetics

**DOI:** 10.1101/2024.11.17.624006

**Authors:** Mafalda Abrantes, Tiago Pereira, Patrícia Silva, Margarida Falcão, Jérôme Borme, Pedro Alpuim, Luis Jacinto

## Abstract

Optogenetics is a widely used tool to dissect neural circuits with optical stimulation, but it requires that light is delivered to photosensitive neurons inside the brain. Implantable neural probes with microscale LEDs (μLEDs) are an emerging approach to delivering light to the brain with superior light output control. However, approaches to integrate μLEDs in neural probes depend on complex fabrication processes. Here, we developed an implantable small form factor neural probe that integrates highly efficient commercial flip chip μLEDs using only standard lithography processes in silicon and a custom automated LED mounting approach with custom 3D-printed tools on a pick-and-place machine. The probe has a cross-sectional area under 0.013 mm^2^ but can output up to 2.5 mW of optical power with an irradiance of 175 mW/mm^2^. Due to the high plug efficiency of the LED, the neural probe can perform stimulation protocols up to 20 Hz and 80% duty cycles without surpassing estimated hotspot temperature elevations above 1 ºC. The neural probes were validated in vivo, with brain activity in the motor cortex of transgenic mice being reliably modulated by pulsed light emitted from the probe.

## 1. Introduction

Optogenetics is a powerful tool to modulate neuronal circuits with optical stimulation, and it has been widely used to dissect functional connections in the brains of preclinical animal models [1]–[3]. Through the genetic modification of neuronal cells under the control of specific promoters to express light-gated channels and pumps, optogenetics allows selective excitation and inhibition of neuronal circuits with millisecond resolution using different light wavelengths [4], [5]. This has allowed researchers to map functional connections between different cell types in brain circuits or between different brain areas, as well as to highlight potential cellular targets for brain disorders [3], [6], [7]. Initial approaches to deliver light inside the brain to activate photosensitive cells resorted to implantable optical fibers connected to an external light source such as a laser or an LED [8]. These systems were widely disseminated as they are easily scalable, and optical fiber implants are simple to fabricate. Over the past decade, advances in optoelectronics, photonics, and microfabrication processes led to alternative approaches in which the light source could be implanted inside the brain. [9]–[11]. This can be achieved, for example, by the integration of microscale LEDs (μLEDs) in micron-size implantable neural probes using silicon-based MEMS fabrication techniques similar to the ones already used for developing neural probes for electrophysiological recordings [9], [12]. Neural probes with μLEDs allow direct light output control with low operational currents and voltages, facilitating multiplexed array designs, wireless operation, and integration in multifunctional devices [12], [13], and open new pathways for optical interrogation of neural circuits.

The integration of μLEDs in neural probes for optogenetics can follow different routes, each with advantages and disadvantages. One route resorts to monolithic integration by fabricating the device directly on sapphire or silicon epitaxial wafers with gallium nitride (GaN) [14]–[16]. This option offers limited substrate choices and can bring additional compromises regarding materials and processes [9], [17]. Because the entire wafer is used for device fabrication, a cumbersome thinning step is often necessary to reduce the probe’s final implant footprint [14], [16]. Another route consists of epitaxial lift-off from wafers with GaN followed by transfer printing to a target substrate such as silicon [18] or polyimide [19], [20]. This approach retains the optoelectronic performance of μLEDs grown on the epitaxial wafers and introduces fabrication flexibility of the final device, including in the choice of substrates that can then be released to create smaller cross-section or flexible neural probes [19], [20]. However, the transfer and bonding steps to the target wafer can be complex, requiring temporary handling wafers or assistive transfer support materials and post-transfer processing steps. Both routes can produce very thin μLEDs (< 5 μm) integrated into reduced thickness neural probes (15-120 μm) but typically suffer from low wall-plug efficiency (< 1-2%) and low optical power and irradiance.

An alternative route is the integration of commercially available small-footprint flip chip μLEDs, that were produced for other applications, through micrometric positioning and bonding to appropriate bonding pads fabricated on the probe’s substrate. But placing and effectively bonding and passivating the chips can be challenging due to their small size and the reduced dimensions of the implantable shanks in the neural probes. Previous approaches have mainly used large area μLEDs (> 0.06 mm^2^) and thicker substrates (up to 700 μm) that can withstand the post-fabrication mounting and bonding processes to overcome such difficulties [21]–[24]. The result is neural probes with large cross-sectional areas, potentially leading to more tissue damage upon implantation and poor performance.

In addition to the implant footprint, another concern with implantable neural probes with μLEDs is that a large portion of the input power is converted into heat, not optical power [25]. Temperature elevations in brain tissue can have modulatory effects on neuronal cell activity or even cause tissue damage [1], [25], [26]. Although there are no official guidelines specific to brain implants, regulations limit surface temperature rises of implantable medical devices to 2 ºC. To stay on the conservative side, there is acceptance that μLED neural probes should not increase brain tissue temperature by more than 1 ºC during operation [25]. Due to the low wall-plug efficiency of μLEDs used in neural probes and the necessity of using higher voltages to achieve reliable turn-on currents, most neural probes with μLEDs are limited to operation with either low driving currents or low frequency and duty-cycle stimulation protocols to avoid unintended temperature elevations above the 1 ºC limit.

Here, we report on an implantable silicon-based neural probe that integrates highly efficient small-sized commercial bare die μLEDs on thin shanks with a small final cross-section and high optical output power. A custom method was developed to mount, bond, and passivate the μLEDs on 15 μm-thin probe shanks using custom-printed 3D tools on a pick-and-place machine. The final cross-sectional area of the probe is only 0.013 mm^2^, which is significantly smaller than that of the smallest optical fiber implants used in mice. The probe can output up to 2.3 mW of optical power with an irradiance of 175 mW/mm^2^, with approximately 15% plug-efficiency while maintaining hotspot temperature elevation below 1 ºC for frequencies up to 20 Hz and duty cycles below 80%. The probe was validated in vivo by reliably driving opto-evoked activity in the motor cortex of transgenic mice.

## 2. Methods

### 2.1 Neural probe fabrication

A standard silicon-on-insulator (SOI) wafer (15 μm silicon (Si) / 2 μm silicon dioxide (SiO_2_) / 650 μm Si / 2 μm SiO_2_) (Silicon Valley Microelectronics) was used (Figure 1a). The fabrication process started with the sputtering of an alumina insulator film (Al_2_O_3_, 100 nm) on the front side of the wafer, and the deposition of 1500 nm of SiO_2_ on the back side by plasma-enhanced chemical vapor deposition (PECVD) (Figure 1b,c). Then, a 5 nm layer of chromium (Cr), as adhesion for the gold (Au) contact layer, and 200 nm of Au were sputtered on the front side (Figure 1d). Probe contacts, pads, and conductive lines were patterned by photolithography and etched by ion beam (Figure 1e,f). On the back side, a SiO_2_ hard mask was patterned by reactive ion etching to define the regions that would later be etched up to the device layer (Figure 1g,h). A layer of 200 nm of Al_2_O_3_ was deposited on the front side by atomic layer deposition (ALD) for device passivation, and probe contacts and pads were exposed by ion milling (Figure 1i,j,k). The outline of each probe was defined by photolithography, and the wafer’s Si device layer (15 μm) was etched by deep reactive ion etching (DRIE) (Figure 1l.m). Then, the back side of the wafer (Si 650 μm) under the probe’s shank was etched with the same DRIE process until the buried 2 μm SiO_2_ layer was reached (Figure 1n). Finally, the buried oxide layer of the neural probe shank was etched by hydrogen fluoride (HF) vapor (Figure 1o,p). Probes were released from the wafer by breaking two thin silicon bridges that connected the probes’ base to the wafer.

**Figure 1.**
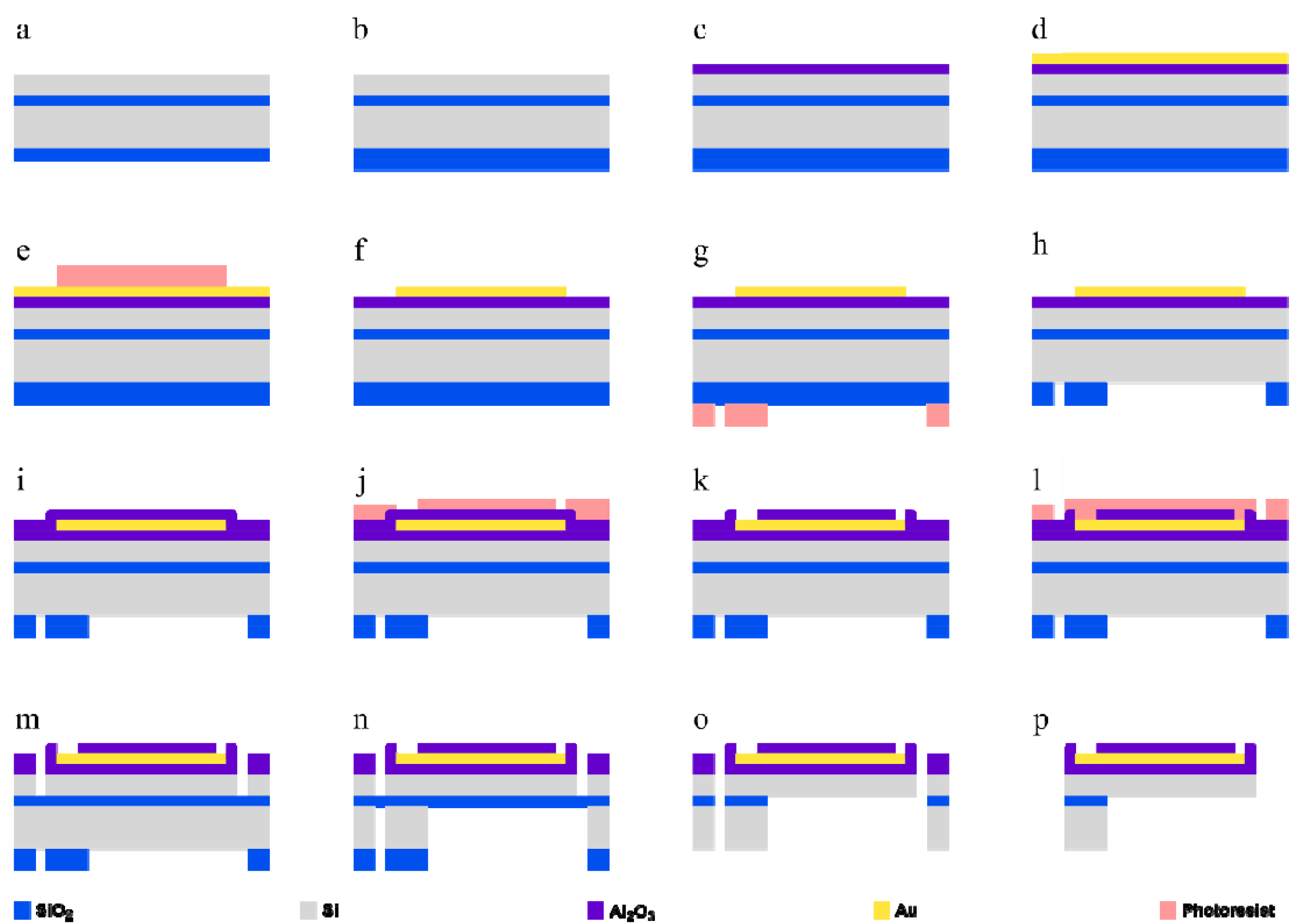
Neural probe fabrication process. (a) Silicon-on-insulator (SOI) wafer. (b) Back-side silicon oxide (SiO_2_) layer deposition. (c) Top-side alumina (Al_2_O_3_) layer deposition. (d-f) Gold (Au) deposition and patterning of interconnect lines and bonding pads. (g-h) Back-side SiO_2_ patterning. (i-k). Top-side Al_2_O_3_ passivation and patterning of bonding and connector pads. (l-m) Top-side etching to define probe layout. (n) Backside etching to define probe layout and remove silicon (Si) from under the neural probe shank. (o) SiO_2_ etching to release device layer. (p) Final released neural probe. NB: not to scale.

### 2.2 μLED integration in neural probe

μLEDs (UB06FP2, Light Avenue) were mounted on the neural probe shank by a pick-and-place system (Fineplacer Sigma, Finetech) with custom-designed holders and tips. A specially designed holder for handling the probe during the process was printed on a stereolithography resin 3D printer (Form 3+, Formlabs) and cured by ultra-violet (UV) light exposure (in Form Cure, Formlabs). Double-sided thermal-release tape (TRT) was taped to the holders to attach the probe’s shanks. Probes were inserted into the holders with the help of tweezers and, after vacuum fixation in the pick-and-place working stage, these were gently pressed (0.02 N) against the TRT with a custom 3D printed tip tool (with a cylinder tip of 500 μm diameter). Conductive adhesive (Ablestick ABP 8037 TI, Loctite) was stamped on each probe’s bonding pad with another stamping tool (150 μm diameter semisphere DAUB tip, Small Precision Tools). Then, the μLEDs were positioned on the pads with a pick-and-place vacuum tool (2-sided Inverted Channel Die Collect, Small Precision Tools) and baked in a furnace to cure the conductive adhesive with a 30 minutes temperature ramp between 25º C and 160 ºC, followed by 45 minutes at 160 ºC. After cooling to room temperature, the μLEDs on the probes’ shanks were passivated with a thin 5-10 μm layer of acrylic-based transparent coating for optical applications (8424 UV, IQ-BOND) with a custom pick-and-place 3D printed tool. Finally, the coating was cured with UV-light exposure for 5 s.

### 2.3 μLED neural probe packaging

The μLED probes were packaged with a custom-designed PCB (5.9 × 6 × 0.8 mm) with electroless nickel immersion gold (ENIG) finishing. Probe connector pads were wire-bonded to PCB pads with 25 μm gold wire in a wire bonder (HB16, TPT), and a connector (853-93-100-10-001000, Mill-Max Mfg. Corp.) was soldered to the PCB.

### 2.4 Electrical and optical characterization

I-V characterization was performed with a source meter unit (2400 SourceMeter, Keithley) connected to the probe’s PCBs with tungsten probes. Voltage sweeps from 2.5 V to 3.2 V were applied with 0.1 V increments, with current limited to 6 mA. Optical output was measured in the same setup by placing a photodiode power sensor (S121C, Thorlabs) above the μLED at approximately 1 mm and connected to a power meter (PM101, Thorlabs). One μLED neural probe was immersed in phosphate-buffered saline (PBS) solution for 12 days, and the current was measured daily to a forward bias of 2.8 V to assess the passivation coating efficiency.

### 2.5 Thermal modeling

Heat transfer simulations were modelled in COMSOL Multiphysics (COMSOL Inc) using the Heat Transfer module. The model consisted of a probe with a body assumed to be silicon (density ρ = 2329 kg m^−3^, heat capacity CP = 700 J kg^−1^ K^−1^, thermal conductivity k = 130 W m^−1^ K^−1^) and with the same geometry as the fabricated neural probes. At the tip of the probe’s shank, two blocks with an area of 89 x 150 μm^2^ were stacked to mimic the μLED geometry. The bottom and top blocks were assumed to be SiC (density ρ = 3216 kg m^−3^, heat capacity CP = 490 J kg^−1^ K^−1^, thermal conductivity k = 690 W m^−1^ K^−1^) and sapphire (density ρ = 3980 kg m^−3^, heat capacity CP = 800 J kg^−1^ K^−1^, thermal conductivity k = 45 W m^−1^ K^−1^), with 30 μm and 50 μm thickness, respectively, adding up to the 80 μm μLED’s total thickness. A thin 10 μm thick layer of PMMA (density ρ = 1200 kg m^−3^, heat capacity CP = 1446 J kg^−1^ K^−1^, thermal conductivity k = 0.193 W m^−1^ K^−1^) surrounding the two μLED blocks was modelled to simulate the transparent acrylic-based coating. The shank was assumed to be inside brain tissue (density ρ = 1040 kg m^−3^, heat capacity CP = 3650 J kg^−1^ K^−1^, thermal conductivity k = 0.527 W m^−1^ K^−1^) with the bulk of the probe being in air (COMSOL’s model). Outside and brain temperature were initially defined at 21°C and 37°C, respectively. Heating was assumed to originate from the walls of the SiC block and the top surface of the probe’s shank and modelled by applying Boundary Heat Sources to its faces. A time-dependent simulation was run in 1 ms time intervals from 0 to 1 seconds. Output power was assumed to be due to Joule Heating in the probe’s shank (P = RI^2^) and the difference of input electrical power and output optical power at the μLED (P = V_μLED_I – P_optical_). Tissue temperature for graphical representations was considered as the average tissue temperature in a 30 μm line from μLED coating surface.

### 2.5 In vivo testing

A custom tetrode device [27] carrying four nichrome tetrodes was used for electrophysiological recordings. Briefly, tetrodes were prepared by twisting four 12.5 μm nichrome wires (Kanthal) with Twister3 [28] and mounted on the tetrode device with UV-cure epoxy. Following tetrodes connection to the pads with gold pins (Neuralynx), the tips of the tetrodes were cut and gold-plated as in [27]. The μLED neural probe connected to the PCB package was then glued to the tetrode device with UV-cure epoxy so that the μLED was facing the tetrodes’ tips at a distance lower than 1 mm.

One Emx1-Cre:Ai27D male mouse was used for in vivo tests. Mice were obtained by crossing homozygous Emx1-IRES-cre mice with homozygous Ai27D mice (JAX stocks #005628, #012567) [29], [30]. The animal was anesthetized by intraperitoneal injection of ketamine (75 mg/Kg) and medetomidine (1 mg/Kg) mix and securely placed in a stereotaxic frame (World Precision Instruments). The tetrodes and μLED neural probe assembly was lowered into the motor cortex (1.7 mm AP and 1.0 mm ML from bregma) to a depth of 0.9 mm (DV) from the brain surface. A stainless-steel screw at the back of the skull served as ground. Extracellular neuronal activity was acquired at 30 kS/s with a headstage (RHD 32ch, Intan) connected to the tetrode device and an Open Ephys acquisition system [31], and filtered between 0.3 and 6 kHz. To evoke neuronal activity from motor cortex neurons expressing channelrhodopsin, the μLED neural probe was powered by two AA batteries at 2.7 V and controlled by an Arduino UNO (rev3, Arduino) to deliver 10 Hz light stimulation pulses (80% duty cycle). The inter-trial interval was 30 s. The Arduino also sent a 3.3 V TTL pulse during stimulation to the acquisition system for synchronization with the electrophysiological recordings. Evoked electrophysiological responses by optical stimulation were analyzed with custom matlab code. Spikes were detected if crossing an amplitude threshold five times higher than the signal’s root mean square (RMS). Detected spikes were averaged in 200 ms bins for the peristimulus time histogram (PSTH).

## 3. Results and discussion

### 3.1 Probe design and fabrication

A small cross-sectional area neural probe with an integrated flip chip μLED that provides high optical power at low driving power was developed in this work for in vivo optogenetics applications. Neural probes were fabricated with standard photolithographic processes on a silicon-on-insulator (SOI) wafer (Figure 2a,b), and the μLED chip was then transferred to the released probes post-fabrication. A SOI wafer was chosen because it allows precise etching of its bottom side to create neural probes with reduced thickness implantable shanks (device layer) and a connector section at full-wafer thickness for safe handling [32], [33]. Probes of 3 and 6 mm lengths were fabricated to allow targeting of different cortical and subcortical brain areas in both rats and mice (Figure 2c). The implantable portion of each probe, regardless of length, consists of a 99 μm wide and 15 μm thick shank with two gold 44 × 33 μm^2^ bonding pads with a vertical pitch of 100 μm near the tip for posterior μLED integration (Figure 3d). Both pads are connected to 40 μm wide gold interconnect lines (200 nm thick), with a lateral separation of 7.5 μm, extending to the connector section of the probe. The width of the shank was primarily determined by the width of the μLED to be mounted (89 × 150 μm), and gold interconnect lines were lithographically patterned to occupy most of the available space to reduce line resistance and improve heat dissipation. The connector section of the probe remained at wafer thickness (670 μm) and included two 990 × 490 μm^2^ gold contact pads that were used to wire-bond the probe to a custom-designed PCB. The line resistance between μLED bonding pads and connector contact pads was calculated to be 25 and 50 Ω for the 3 and 6 μm length probes, respectively. The width of the interconnect lines could be reduced to include additional μLEDs in the shank, but to retain similar line resistances, the thickness of the deposited gold would have to be increased. The design of probes with multiple shanks to simultaneously target different brain areas could also be realized without changes to the overall fabrication process.

**Figure 2.**
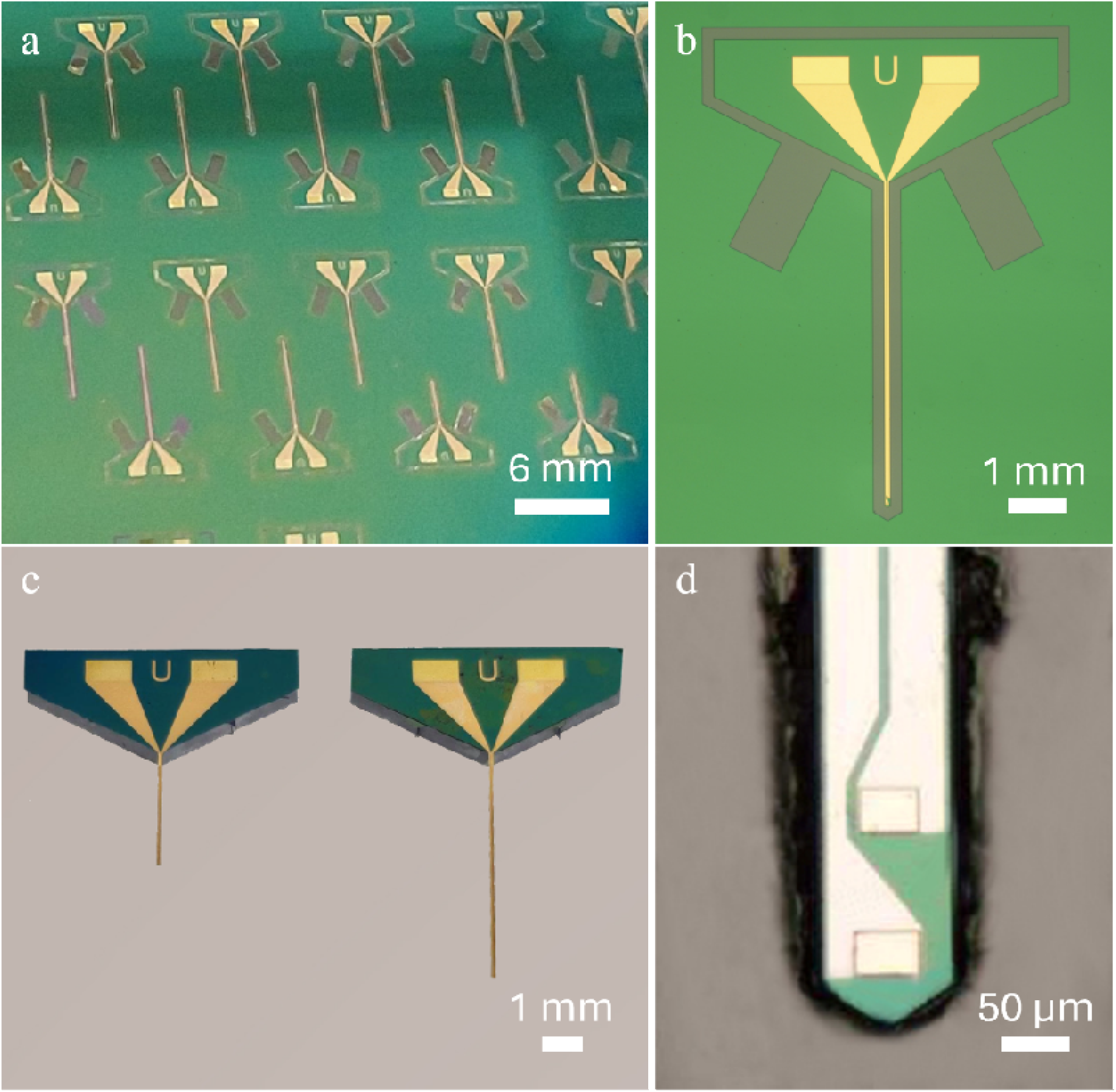
Fabricated neural probes. (a) Detail of SOI wafer with neural probes during the fabrication process. (b) Optical microscopy photograph of neural probe on wafer. (c) Two neural probes released from the wafer, with 3 and 6 mm implantable shanks (left and right, respectively), ready for the μLED integration process. (d) Optical microscopy photograph of neural probe shank’s tip on wafer before the final backside etch for release, showing the wide interconnect lines and the two bonding pads for μLED mounting.

**Figure 3.**
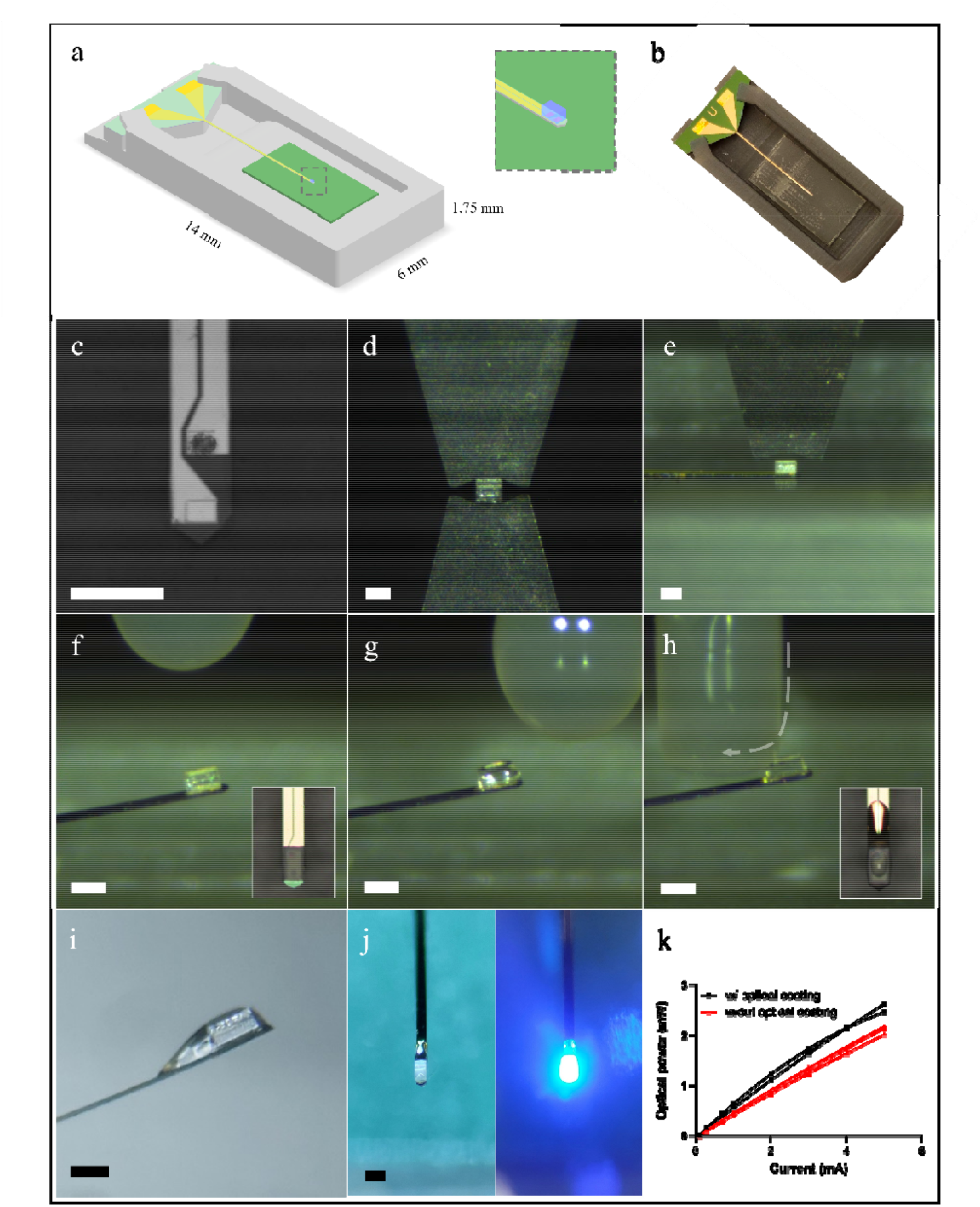
μLED integration on neural probe. (a) Schematic of custom 3D-printed probe holder with an inserted probe. The green patch under the probe’s shank corresponds to the termal release tape (TRT). (Inset) Close-up schematic of a bonded μLED in the shank’s tip. (b) Photograph of a 6 mm neural probe on a 3D-printed holder with TRT. (c) Shank’s tip with the top bonding pad covered in conductive glue. (d) μLED picking process using the vacuum tip. (e) μLED placement on the shank’s tip supported by the TRT on the probe holder. (f) Neural probe shank with bonded μLED, released from the TRT after curing. (Inset) Optical microscopy photograph with top view of the μLED in the probe after curing. (g) μLED after application of the optical coating applied with the custom 3D-printed tool. (h) Wiping excess optical coating of the μLED’s top. Grey arrow denotes the tip’s movement direction. (Inset) Optical microscopy photograph with top view of the μLED after the wiping process. (i) μLED profile photograph after the integration process. The optical coating creates a tail extending away from the tip that smoothens the probe’s tip topography. (j) Photograph with top view of the finalized probe in a OFF (left) and ON (right) states. (k) Measured optical power output of μLEDs with (black lines) and without (red lines) optical coating. Scale in (c),(d),(e),(f),(h),(i) and (j) is 150 μm.

### 3.2 μLED integration

Integrating bare μLED chips in neural probes allows using highly efficient μLEDs but typically requires thick substrates to withstand the mounting and bonding processes primarily performed post-fabrication. In contrast to previous approaches that led to thick implantable probes (up to 700 μm) [21]–[23], a novel method using custom 3D-printed holders and tips for a pick-and-place system was developed to integrate bare μLED chips with reduced dimensions (89 × 150 × 80 μm^3^) in 15 μm thick suspended silicon neural probe shanks. When compared with other bare μLED chips previously used in neural probes, such as the TR2227 (220 × 270 × 50 μm^3^, CREE) [21], [22], [24], the chosen LED for our probes has a considerably smaller footprint albeit being 30 μm thicker. The integration of the μLEDs on the neural probes had to be realized after the final fabrication step, that includes a hydrogen fluoride (HF) etch process to remove the buried oxide layer at the bottom of the shanks, because the LED chips could not withstand this process.

Custom 3D-printed holders were designed to hold the released neural probes tightly in position during the μLED bonding process (Figure 3a,b), allowing for an accurate and reproducible process. The 3D printed pieces were designed to have two distinct areas that could accommodate the thickness differences between the probes’ shanks (15 μm) and their connector portion (670 μm). Double-sided thermal release tape (TRT) was used in the holders to guarantee that the probe’s shank could not move laterally and provide cushioning during the subsequent stamping and bonding procedures (Figure 3b). Different pick-and-place tips, including custom-designed ones, were used to sequentially press the probe’s shank against the TRT, stamp conductive glue on the shank’s bonding pads, and position and press the μLED on top of the bonding pads (Figure 3c,d,e). Following conductive glue cure in a furnace for 75 mins, where the probes were also released from the TRT, a custom 3D-printed tool was used in the pick-and-place system to passivate the μLED with a thin layer of a UV-cured transparent optical coating (Figure 3f,g). The custom-designed tool creates a coating fluid dome that envelops the μLED and guarantees both the coating of the LED and underfilling between the LED and the neural probe. The tool also allows the final wiping of any excess coating on top of the μLED for a uniform top layer that is only 5-10 μm thick (Figure 3h). The wiping process also extends upward along the shank, away from the μLED, to create a coating slope that reduces the sharp topography change between the shank and the μLED (Figure 3i). This approach is simultaneously more reproducible and more effective in maintaining a reduced probe thickness than, for example, dip coating, which has been used in previous μLED neural probes [22] but can double their total thickness. An exploratory preliminary test also showed that hexamethyldisilazane (HMDS) vapor priming, which is a standard process in microfabrication for adhesion promotion between substrates and photoresists, increases the wettability of the μLEDs and improves coating distribution and underfilling with the described passivation process. The full μLED mounting and passivation process had a yield of approximately 80%, with sources of error arising from mishandling the probes, misalignment of the μLEDs and the contact pads, and insufficient optical coating being applied. The necessity of handling the released probes and inserting them into the custom 3D holder led to some probes being lost due to human error. During stamping of conductive epoxy there was a risk of short-circuiting the two gold pads either by overapplying glue or by sub-optimal alignment of the stamping tool and substrate, but these occurred rarely. The suboptimal in-plane and out-of-place alignment of μLED and stamped gold pads could also lead a poor connection/adhesion, which resulted in the μLED being released from the probe during the curing of the conductive epoxy or in an open circuit. This error typically arised from picking up the μLED in a tilted position or miscalculating the alignment of the μLED holder and the probe and holder. Although these sources of errors became very infrequent once the user had gained experience with the process, typically after 20 probes, some process automation could be implemented to increase both its yield and scalability. Designing a wafer-sized custom holder, instead of individual probe holders, with pre-defined positions for the probes on the wafer would eliminate the need for human handling of the fragile probes. Furthermore, this could allow for computer vision pattern recognition to be adopted both for stamping the conductive glue and picking and placing the μLEDs at the wafer level. This type of pick-and-place automation is well established in industry for assembling micro-components in PCBs, including advanced systems capable of a higher number of degrees of freedom that could be used to prevent out-of-plane misalignments between μLED and bonding pads. The use of HMDS vapor priming, as described above, could also contribute to improving the application of the coating. The scale-up of this fabrication process extends to any type of available μLED that could fit the probe or to which the probe design could be adapted. The versatility of the process means that if smaller footprint μLEDs become available, smaller probes with similar characteristics could be fabricated with minimal process modifications.

The specific optical coating was chosen because of its optical transparency between 230-500 nm, which encompasses the emission wavelength of the used μLED (465 nm) (Figure 3j), and its reported humidity resistance. To confirm that the optical coating did not reduce the optical output of the μLED, optical power was measured before and after the coating application. An increase in μLEDs optical power after passivation was observed (Figure 3k). This was most likely due to a reduction of the solid angle emission of the LEDs, considering a planar photodetector and not an integrating sphere was used to measure optical power (if using the latter, output power between coated and uncoated LEDs was expected to be similar). The final cross-sectional area of the neural probe, considering the substrate, μLED, and coating, did not exceed 0.013 mm^2^, which is not only smaller than in most μLED probes, regardless if they were monolithically fabricated or not, but also smaller than the smallest 100 μm optical fiber implants used in mice implants (having a footprint of approximately 0.050 mm^2^ considering the core, cladding, and coating). Table 1 shows a comparison of our neural probe’s characteristics with those that have been previously published.

**Table 1.** Comparison of different implantable neural probes with μLEDs for brain optogenetics.

| μLED | μLED dimensions (w x l x h) (μm) | μLED emission peak (nm) | Probe shank dimensions (w x h) (μm) | Cross-sectional area ($\text{mm}^2$) | μLEDs per shank | Probe substrate | Safe* max working current (mA) | Optical power at safe* current (mW) | Irradiance at safe* current ($\text{mW}/\text{mm}^2$) | Peak Wall-plug efficiency (%) | Temperature below 1 °C | In vivo testing | Ref |
| --- | --- | --- | --- | --- | --- | --- | --- | --- | --- | --- | --- | --- | --- |
| Commercial flip chip LED | 89 x 150 x 80 | 465 | 99 x 105 | 0.014 | 1 | Si | 0.75 | 0.5 | 40 | 14.7 | Below 20% duty cycles for currents up to 2.3 mA; or below 80% duty cycles for 0.75 mA | Mice, cortex M1 | This work |
| GaN-Si | 10 x 15 x 0.5 | 460 | 70 x 30 | 0.0021 | 3 | Si | 0.013 | 0.0003 | 5 | 0.8 | Below 3.4 V | Mice, hippocampus CA1 | [14] |
| GaN-Si | 25 (diameter) | 450 | 100 x 40 | 0.0040 | 16 | Si | 5 | 0.06 | 125 | 0.8 | Below 10% duty cycles | Mice, cortex | [15] |
| GaN-Si | 310 x 41 | 450 | 220 x 100 | 0.022 | 3 | Si | 3.5 | 0.06 | 1 | 0.4 | - | - | [39] |
| GaN-Si | 22 x 22 x 5 | 445 | 800 x 15 | 0.012 | 32 | Parylene C | 0.0085 | 0.0013 | - | 0.5 | Max 5 ms pulse | - | [16] |
| GaN-Si | 50 (diameter) | 460 | 300 x 300 | 0.090 | 5 | Si | 1 | - | 15 | 1 | Below 1 mA currents | Mice, hippocampus | [38] |
| GaN-Sa transferred to PI | 50 x 50 x 6.45 | - | 300 x 20 | 0.0060 | 4 | PI | - | - | 40 | 1.5 | Below 20 Hz | Mice, VTA | [19] |
| GaN-Sa transferred to PI | 125 x 180 | 480 / 630 | 320 x 120 | 0.038 | 1 | PI | 3 / 10 | - | 60 / 45 | 8.0 / 4.0 | Below 3 mA currents / Below 30 Hz up to 10 mA | Mice, VTA | [20] |
| GaN-Sa transferred to Si | 100 (diameter) | 455 | 150 x 65 | - | 10 | Si | 5 | 0.450 | 56 | 2.0 | Below 3 mA currents and 500 ms pulses | Mice, cortex S1 | [18] |
| GaN-Sa | 250 x 490 | 444 | 400 x 200 | 0.080 | 1 | Sa | 35 | - | 5 | - | - | - | [34] |
| GaN-Sa | 40 (diameter) | 450 | 80 x 101 | 0.0081 | 5 | Sa | 0.1 | - | 10 | - | Below 50 Hz | - | [37] |
| Commercial flip chip LED | 220 x 270 x 50 | 470 | 350 x 130 | 0.045 | 1 | PI | 5 | - | 50 | - | Below 60% duty cycles | Mic, Nac / VTA | [22] |
| Commercial flip chip LED | 220 x 270 x 50 | 460 | 250 x 60 | 0.016 | 3 | PI | 10 | 0.5 | 10 | 1.7 | - | Mice, nRT | [24] |
| Commercial flip chip LED | 220 x 270 x 50 | 465 / 540 | 650 x 700 | 0.45 | 2 | PDMS | - | - | 12 | - | Below 70% duty cycles | Mice, Nac | [21] |
| Commercial flip chip LED | 290 x 550 x 100 | 455 | 900 x 500 | 0.45 | 1 | PCD | - | - | 1.55 | - | Below 1 Hz and 100 ms pulses | Rat, V1 | [23] |

\*Safe working current or voltage were defined as the current or voltage values that do not lead to increases over 1 °C in probe hotspot temperature or brain tissue with the most conservative driving parameters; if temperature assessment was not provided, the maximum working current is reported (when available). Abbreviations: CA1 – hippocampus cornus ammonis 1; GaN – gallium nitride; M1 – primary motor cortex; nRT: nucleus reticularis thalami; PCD – polycrystalline diamond; PDMS – Polydimethylsiloxane; PI – polyimide; S1 – primary somatosensory cortex; Sa – sapphire; Si – silicon; VTA – ventral tegmental area

### 3.3 Electro-optical and thermal characterization

The electrical and optical properties of the fabricated μLED neural probes were evaluated with a source meter and a photodetector on a probe station. Representative I-V curves and radiant flux as a function of current were obtained for the probes with μLEDs (Figure 4a,b,c). Turn-on-voltage was observed at 2.6 V. The measured optical output power and calculated irradiance at driving voltages between 2.6 and 3.2 V (0.3 to 5 mA) ranged between 0.15 and 2.5 mW and 10 and 175 mW/mm^2^, respectively (Figure 4b,c). Although only 3 and 4 probes were tested to construct the representative I-V curves and radiant flux curves, respectively, the electrical and optical measurements presented very low variability across probes as can be observed by the low standard deviations (Figure 4b,c) indicating high μLED operational stability and reproducibility of the fabrication method. The irradiances obtained were significantly higher than those reported for previous μLED-based probes at the same driving voltages/currents (Table 1). The high irradiance in our probes is due to the small emission area of the LED chip (89 × 150 μm^2^) and its high wall-plug efficiency (WPE). The observed WPE of 14.6% is significantly higher than those previously reported for any μLED neural probe (Table 1), allowing our probe to output at least two orders of magnitude more optical power than most previous probes with similar currents (and with many also requiring higher operating voltages) [15], [18], [20], [22], [24], [34].

**Figure 4.**
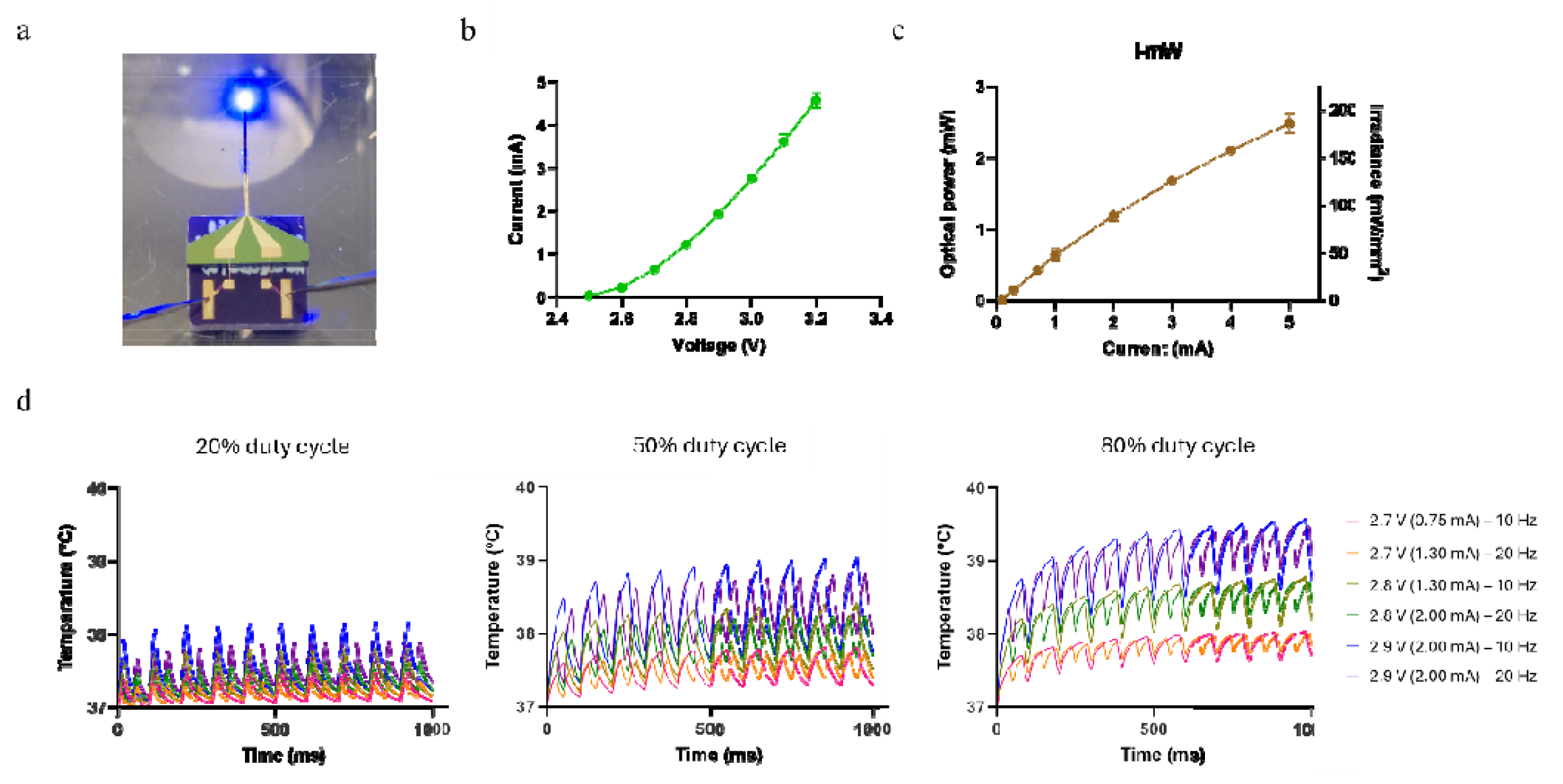
Opto-electrical and thermal modeling of neural probes with μLED. (a) Photograph of probe wire-bonded to the PCB and driven by tungsten needles from probe station. (b) Representative I-V curve (n=4, error bars are s.d.). (c) Representative optical power and irradiance of neural probe with μLED as a function of driving current (n=3, error bars are s.d.). (d) Simulation of brain tissue temperature modulation, based on COMSOL thermal modeling, for μLED pulsed operation (10 and 20 Hz) for different driving voltages (2.7, 2.8, and 2.9 V) and duty cycles (20%, 50%, and 80%; left, middle, and right panels, respectively) during 1 second.

LEDs are never 100% efficient, and a significant part of the driving power is converted to heat not contributing to the optical output [25]. Because many neuronal circuit processes are temperature dependent, it is crucial to estimate or assess brain tissue temperature elevations due to hotspot temperature in the μLED during optical stimulation with different stimulation parameters typically used in optogenetics experiments [25], [26]. To evaluate the potential heating of brain tissue due to μLED probe operation, a model was developed in COMSOL Multiphysics (COMSOL Inc.) using the heat transfer module. The probe geometry was modeled with the exact dimensions of the fabricated neural probes and the μLED and the optical coating were considered in the model. The μLED heating was assumed to be due to the dissipated power corresponding to the difference in input electrical power and output optical power. The heating of the brain due to optical absorption of the emitted light was assumed to be negligible. Several simulations were ran for all possible combinations of the following stimulation driving parameters: 2.7, 2.8, and 2.9 V (0.75, 1.3, and 2 mA, respectively), 10 and 20 Hz frequency, and 20, 50, and 80% duty cycles (Figure 4d). The observed induced temperature changes in the brain tissue close to the μLED were dependent on the μLED driving parameters, with higher voltages, lower frequencies, and longer duty cycles inducing the highest temperature rises. The longer the μLED was turned on at each driving voltage (i.e. lower frequencies and/or longer duty cycles) the higher the heating, as previously reported for other neural probes [14], [15], [18], [19], [22]. From the tested driving parameters, all possible combinations with 20% duty cycles (except 2.9 V at 10 Hz) lead to brain tissue temperature rises below 1 ºC. With 50% and 80% duty cycles, only voltages below 2.7 V (0.75 mA) did not raise brain tissue temperature above the safe limit of 1 ºC. Using these lower input power settings, an irradiance of approximately 25 mW/mm^2^ is expected, which well above the threshold of 1 mW/mm^2^ necessary for channelrhodopsin activation [35].

Compared with previously reported μLED neural probes, our probe presents lower heating for similar driving power (Table 1). This can be partly explained by the significantly higher WPE of the μLED used and the potential reduction of the hotspot temperature through heat blocking by the 10 μm optical acrylic passivation coating which has low thermal conductivity and can block heat transfer. A modeling study estimated that μLED hotspot temperatures on neural probes can decrease by 50% with only 20 μm of encapsulation [25]. The heating of brain tissue through μLED operation is also likely lower than modeled because of the large heat capacity of brain tissue that prevents significant temperature elevations. Blood flow in the brain’s vasculature works as an active heatsink that contributes to actively modulate brain temperature [36]. Previous studies support that temperature elevations in the tissue will be even lower than those measured or modeled at neural probe’s surface [14], [37], [38]. Heat dissipation through metal interconnect lines, working as temporary heat buffers due to their high thermal conductivity, can also contribute to reduce μLED’s hotspot temperature in neural probes [15], [37] [16], [25]. Previous reports showed that relevant heat dissipation in μLED probes can occur through metal lines and can even lead to considerable hotspot temperature elevations in the exposed electrode sites of probes combining capacitive metal electrodes with μLEDs [16], [25]. However, the gold interconnect lines in our probe were not included in the simulation model because adding a nanometric layer would increase the the required computing power significantly. Therefore, it cannot be discard that a proportion of heat dissipation likely occurred through the interconnect lines due to the high thermal conductivity of gold potentially leading to lower μLED’s hotspot temperature during operation.

### 3.4 In vivo testing

Before in vivo testing in mice, the longevity of the μLED optical coating encapsulation was assessed. The shank of a neural probe was immersed in phosphate-buffered saline (PBS) continuously for 12 days, and current measurements at 2.8 V were taken every 24h (Figure 5a). A current of approximately 1.30 mA was measured every day (Figure 5a), confirming that the regular operation of the probe was maintained even in continuous immersion in liquid solution. The process of immersing μLED probes in PBS was repeated with another 5 randomly chosen neural probes for different durations (5-12 days) but current measurements were not taken. Functioning of the probes was only visually confirmed by driving them with different voltages (2.7 to 3.2 V) in PBS at the end of the defined immersion period. All tested probes were fully functional after prolonged immersion.

**Figure 5.**
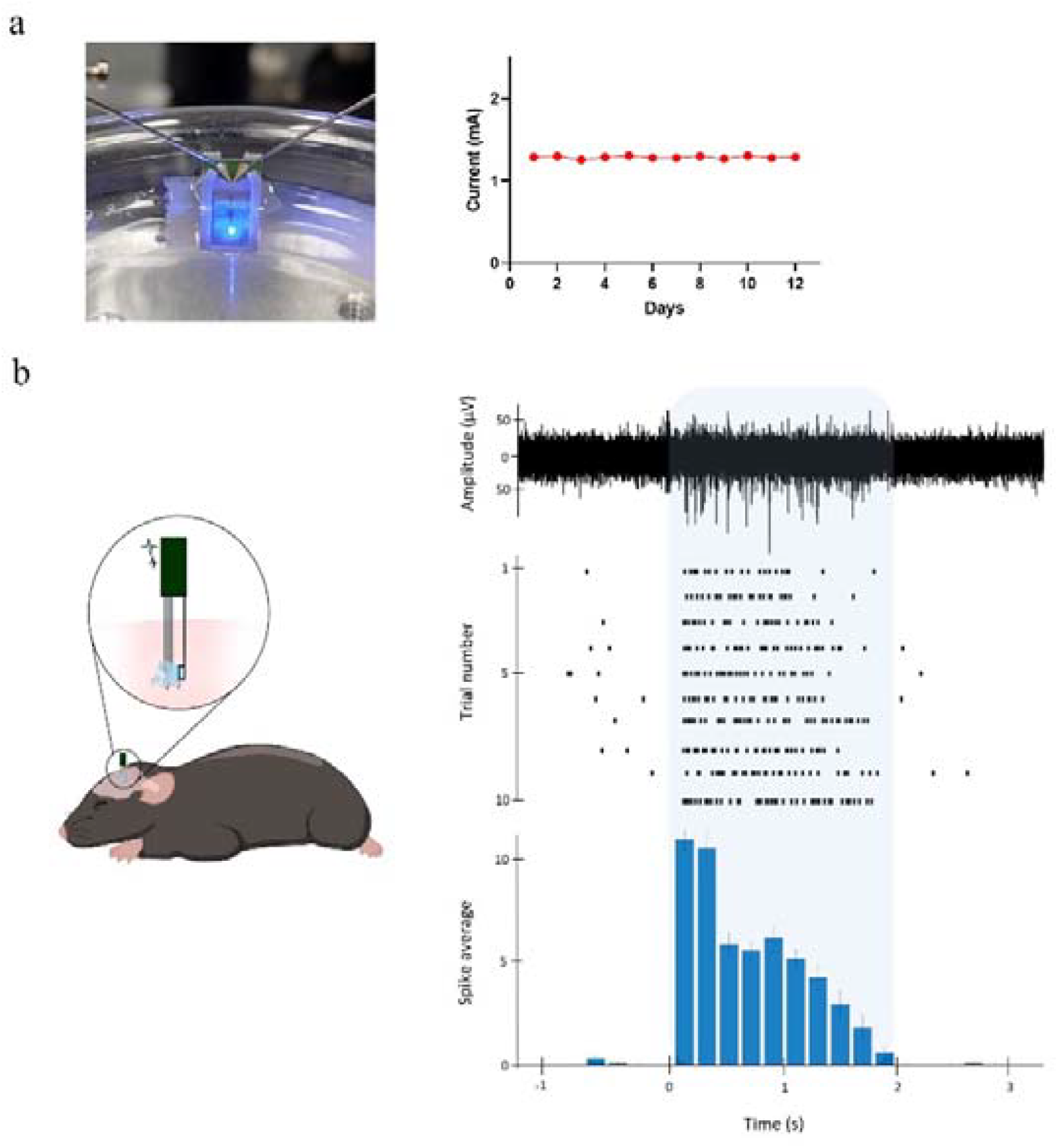
Passivation and in vivo testing. (a) Neural probe submerged in phosphate-buffered saline (PBS) with needles from probe station driving μLED to emit light (left). Current measured daily at 2.8 V during 12 days of continuous immersion in PBS (right). (b) Cartoon of tetrode device and neural probe with μLED assembly implanted in mouse’s motor cortex (M1) for simultaneous electrophysiological recordings and optical stimulation (left). Raw trace (bandpass filtered 300-3000 Hz) of one optical stimulation trial with electrophysiological activity before, during and after optical stimulation (righ, top). Raster plot of spikes detected in 10 optical stimulation trials (right, center). Cumulative peri-stimulus time histogram (PSTH) with total spike count across all optical stimulation trials (right, bottom). Blue shaded area corresponds to optical stimulation period. Bins in PSTH are 200 ms.

To assess the in vivo performance of the μLED neural probe, a custom tetrode device [27] carrying 4 nichrome tetrodes was combined with a μLED probe for simultaneous optical stimulation and electrophysiological activity recordings (Figure 5b). One anesthetized Emx1-cre:Ai27D mouse expressing channelrhodopsin in brain cortical neurons was acutely implanted with the tetrodes-μLED probe assembly in the primary motor cortex (M1). Pulsed brain illumination through the μLED at 2.75 V (approximately 1 mA) reliably induced spiking activity across all trials (Figure 5b). On each trial, the multi-unit activity (MUA) spike rate increased from below 1 Hz to over 25 Hz, confirming strong neuronal activation with optical stimulation. The capability of the neural probe to drive neuronal activity when implanted inside the brain was only tested in one anesthetized animal, and further tests with chronic implants in freely moving mice will be needed to analyze its application in neuroscience behavioral experiments.

## 4. Conclusion

In contrast to previous μLED neural probes with monolithically integrated or transfer-printed LEDs with low quantum efficiency and low output power or commercial bare LED chip bonding on thick substrates, we have designed a novel process to integrate low-cost and highly efficient LED chips in neural probes with reduced thickness (15 um). This approach created small form factor implantable neural probes with a cross-section lower than that found in the smallest optical fibers used for in vivo optogenetic studies and with high optical output power with relatively low operating power/current. In vivo validation with simultaneous electrophysiological recordings confirmed that our neural probe can reliably induce strong spiking activity in neuronal populations expressing channelrhodopsin.

## Acknowledgments

The authors would like to thank Helder Fonseca and José Fernandes of INL for assistance with the HF etching process, as well as all members of the Neurophysiology and Neuroengineering Lab at FMUP and the Alpuim group (former 2D Materials and Devices group) at INL for discussions on probe design and fabrication processes. This work was supported by “la Caixa” Banking Foundation under grant agreement LCF/PR/HR21-00410. MA and MF were supported by Portuguese Foundation for Science and Technology (FCT) doctoral fellowships 2022.14536.BD and 2023.00604.BD, respectively.

## Author contributions

MA and TP designed the probe layout. MA fabricated the probe, with assistance from JB. TP and JB developed the μLED mounting process. PS designed the interface PCBs and performed probe characterization measurement with TP. TP performed thermal modelling simulations. MF and PS performed in vivo testing. PS and LJ analyzed electrophysiological recordings. MA, TP, PS, and LJ wrote the first draft of the manuscript. PA and LJ revised the manuscript, acquired funding, and supervised work. All authors read and approved the final version of the manuscript.

## Statements and Declarations

### Data availability

All data supporting the findings of this study are available in the paper and its Supplementary Information section.

### Ethics approval

All procedures involving animals were conducted in accordance with European Union Directive 2016/63/EU and the Portuguese law on the protection of animals for scientific purposes (DL No 113/2013), and approved by the Portuguese National Authority for Animal Health and the Animal Welfare Committee at FMUP (ORBEA-FMUP).

